# A novel rationale for targeting FXI: Insights from the hemostatic miRNA targetome for emerging anticoagulant strategies

**DOI:** 10.1101/501676

**Authors:** Jamie Nourse, Sven Danckwardt

## Abstract

Therapeutic targeting of blood coagulation is a challenging task as it interferes with the delicate balance of pro- and anticoagulant activities. Anticoagulants are employed in millions of thrombophilic patients worldwide each year. The treatment and prevention of venous thromboembolism has changed drastically with the replacement of traditional anticoagulant vitamin K antagonists by direct oral anticoagulants (DOACs), which selectively target coagulation factors Xa or IIa. However for a growing population with comorbidities satisfying therapeutic options are still lacking and the quest for novel therapeutics continues. Recently targeting factors XI or XII have emerged as new therapeutic strategies. As these factors play important roles in thrombosis, nevertheless are practically functionally dispensable for hemostasis, they may potentially overcome the functional obstacle of treating or preventing thrombosis without affecting hemostasis. Based on the recent elucidation of the hemostatic miRNA targetome, we introduce and discuss a hitherto unrecognized rationale for the therapeutic targeting of factor XI. This is based on mimicking endogenous factor XI expression control by therapeutic delivery of miRNA mimics. We discuss the functional difference between various gene targeting approaches, and propose the hemostatic system to represent an ideal model for assessment of the efficacy and safety of such therapeutic components, ushering in a novel therapeutic era with broad applicability.

## 1. Introduction

Cardiovascular disorders including myocardial infarction, ischemic stroke and venous thromboembolism (VTE) are the leading global cause of mortality with over 17 million deaths annually (Lozano, et al., 2012). The incidence of VTE increases markedly with age, starting in the late 40s, with a dramatic increase occurring at 60 years of age (Silverstein, et al., 1998). Around 30% of patients experience a recurrence within 10 years, underscoring a high demand for therapeutic intervention to treat and prevent VTE (Connors, 2017).

Therapeutic targeting of blood coagulation is an inherently difficult task as it interferes with the delicate balance of pro- and anticoagulant activities. Anticoagulants are employed in millions of thrombophilic patients worldwide each year to limit the otherwise essential function of blood clotting (Wendelboe & Raskob, 2016). However this intervention inevitably leads to an elevated risk of undesired bleeding. While some bleeding events are clinically mild and less critical, others can be life threatening, such as intracranial or abdominal bleeding.

Here we provide insights into the distinctiveness of the hemostatic system and illustrate the challenges and opportunities associated with therapeutic targeting. In addition, we present an interdisciplinary contribution to the exciting discussion about next-generation anticoagulants from the area of RNA biology, and how this might foster the implementation of miRNA therapeutics in other disciplines as well.

## 2. The hemostatic system

Hemostasis is accomplished through a network of processes that include platelets, coagulation, anticoagulant and fibrinolytic pathways, which all support the dynamic equilibrium that provides proper blood flow. Such processes have evolved to maintain the blood in a fluid state under physiologic conditions and to stop bleeding after vascular injury (Borissoff, Spronk, & ten Cate, 2011). Disruption of this well-regulated balance leads to pathologic conditions, such as thrombosis or bleeding.

The response to vascular injury culminates in the formation of a platelet plug, the generation of a fibrin clot, the deposition of white cells in the area of tissue injury, and ultimately the initiation of wound repair. During initiation of blood coagulation, tissue factor (TF), the main trigger of coagulation, is exposed at the site of the vascular lesion (Figure 1). The activation of the TF (extrinsic) pathway results in the formation of a catalytic complex with factor VIIa leading to subsequent activation of factors IX and X. The so-called prothrombinase complex, consisting of activated factors X and V promotes a downstream enzymatic cleavage of prothrombin, which yields small amounts of thrombin. Thrombin is pleiotropic; it acts as a central coagulation enzyme that converts fibrinogen into fibrin but also has a critical role in the activation of platelets and factor XIII to induce fibrin polymerization, a fundamental process in the formation of a stable clot or thrombus. In addition, thrombin plays a crucial part in the amplification and propagation of coagulation by supporting positive-feedback activation of the upstream factors V, VIII and XI. Together with factor VIIIa, factor IXa forms the so-called tenase complex which triggers a burst of additional thrombin that is essential for the formation of sufficient fibrin and the sealing of the vascular defect.

**Figure 1.**
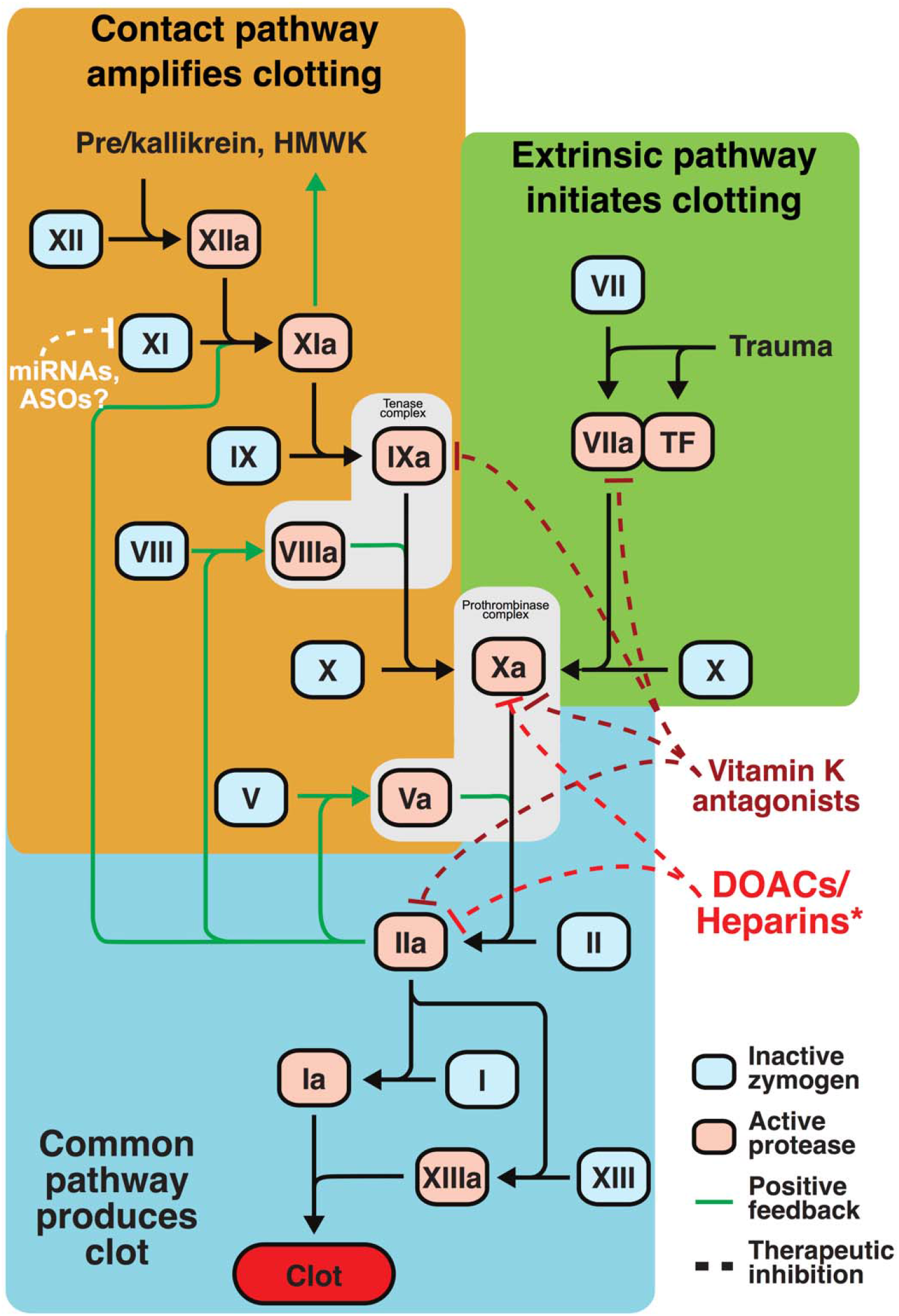
New rational therapeutic targets in the hemostatic system. The hemostatic system is tightly balanced to reduce blood loss by formation of blood clots (hemostasis), while at the same time preventing pathological clot formation (thrombosis). Deficiencies of the intrinsic pathway factors XII and XI are usually not associated with abnormal hemostasis but appear to prevent thrombus formation, qualifying them as attractive therapeutic targets. As FXI is exclusively targeted by miRNAs it provides an attractive avenue for therapeutic targeting (further details see text). miRNA-mediated control of gene expression provides an attractive avenue for therapeutic targeting of factor XI as opposed to current therapies which target the extrinsic and common pathways (such as Vitamin K antagonists, direct oral anticoagulants (DOACS) or antithrombin dependent* (low molecular weight) heparins).

In contrast to the extrinsic pathway, which is essential for protection against bleeding, the contact activation (intrinsic) pathway primarily serves to amplify blood coagulation (Grover & Mackman, 2019) (Figure 1). The exposure of plasma prekallikrein, high-molecular weight kininogen, and factors XI and XII to negatively charged molecules including platelet-derived polyphosphate (Muller, et al., 2009) and collagen (van der Meijden, et al., 2009), results in the conversion of prekallikrein to kallikrein, which activates factor XII. Activated factor XII in turn activates factor XI, and promotes the conversion of additional prekallikrein to kallikrein (see above) which reciprocally amplifies the cascade (Gailani & Broze, 1991; Naito & Fujikawa, 1991). This sequence ultimately leads to the activation of factor IX resulting in proteolytic activation of factor X, culminating in the convergence of both coagulation pathways.

## 3. Current anticoagulants and their limitations

The treatment and prevention of VTE is under a period of rapid change. The introduction of direct oral anticoagulants (DOACs), which specifically target blood coagulation factors Xa (Rivaroxaban, Apixaban, Edoxaban) or IIa (Dabigatran), has changed the management of VTE (Wu & Matijevic-Aleksic, 2005). They are replacing traditional anticoagulants, such as vitamin k antagonists (VKA), which more broadly impair the biosynthesis of factors II, VII, IX and X (Figure 1). DOACs have several advantages; this includes predictable pharmacokinetics, a wider therapeutic window, rapid onset and offset of action, a shorter plasma half-life, fewer food–drug and drug–drug interactions, fixed dosing with - in most cases - no need for laboratory monitoring or dietary discretion, and improved patient satisfaction (Barkun, et al., 2019; Cha, et al., 2017; Y. H. Chan, et al., 2019; Inoue, et al., 2019; Jansson, et al., 2020; Kjerpeseth, et al., 2019). Large clinical trials have demonstrated the superiority of DOAC over VKA in terms of efficacy and safety of VTE treatment (Agnelli, et al., 2013; Bauersachs, et al., 2010; Buller, et al., 2013; Schulman, et al., 2014). An improved benefit/risk profile has also been documented for nonvitamin K anticoagulants regarding the prevention of embolic events and bleeding in atrial fibrillation (Connolly, et al., 2009; Giugliano, et al., 2013; Granger, et al., 2011; M. R. Patel, et al., 2011), and selective Xa inhibition improves the cardiovascular outcome (risk reduction of death from myocardial infarction, or stroke) in patients with recent acute or stable coronary syndrome (Eikelboom, et al., 2017; Mega, et al., 2012).

Nevertheless, despite these improvements, bleeding is still the most frequently reported side effect of DOACs (Eikelboom, et al., 2017; Lanas, et al., 2015; Lanas-Gimeno & Lanas, 2017; B. K. Martinez, Sood, Bunz, & Coleman, 2018; Mega, et al., 2012; van Montfoort & Meijers, 2013; Villines, et al., 2019). Further, selected sub-groups of patients do not show a significant benefit of DOAC with regard to VTE recurrence and bleeding risk (Agnelli, et al., 2015; Prins, et al., 2014; Schulman, et al., 2015). Accumulating studies also report that dabigatran is associated with esophagitis and gastric ulceration, suggesting that the drug may directly injure the gastrointestinal mucosa (Cangemi, et al., 2017; Lanas, 2019; Sostres, et al., 2019). Additionally, satisfying therapeutic options are still lacking for a growing population with co-morbidities who exhibit an increased risk of bleeding under DOACs (Burnett, et al., 2016; Martin & Key, 2016). Apixaban and rivaroxaban are contraindicated in patients with severe renal impairment or advanced liver disease (Abraham, et al., 2015; Alexander, et al., 2019; Andersson, Svanstrom, Lund, Pasternak, & Melbye, 2018; Beyer-Westendorf, et al., 2019; Camm, et al., 2019; Fralick, et al., 2020; S. Y. Lin, et al., 2020; Nissan, et al., 2019; Sakuma, et al., 2019). In patients with mechanical heart valves the use of dabigatran is associated with increased rates of thromboembolic and bleeding complications compared to warfarin, thus showing no benefit and an excess risk (Eikelboom, et al., 2013). As uptake and metabolization of DOACs are P-Glycoprotein- and Cytochrome 3A4-dependent, co-medication with Cytochrome 3A4 and P-Glycoprotein inhibitors (which includes a substantial number of standard pharmaceuticals) can significantly alter DOAC concentrations and increase bleeding risk (Gong & Kim, 2013). Finally DOACs appear to be associated with an increased risk in patients with antiphospholipid syndrome (Pengo, et al., 2018) limiting their use in this critical population with a significant risk of thromboembolic events.

## 4. Contact system as novel target for anticoagulation therapy

The DOAC targets, factor IIa and Xa, are located downstream in the common pathway (Figure 1) and play essential roles in physiological hemostasis. Targeting either one disrupts both intrinsic and extrinsic coagulation pathways, preventing the positive feedback activation of factors XI, VIII and V by factor IIa generation. Consequently, DOACs have a strong impact on the clotting process, and result in an increased bleeding risk (Gale, 2011). Owing mainly to this concern, their clinical application is still restricted and therapeutic monitoring requirement is controversial (Lesko, 2016; Shoeb & Fang, 2013).

In the search for new and safer anticoagulant therapies, bleeding side-effects may be avoided with the selective targeting of coagulation factors that play more important roles in pathological thrombosis than for physiological hemostasis (Plow, Wang, & Simon, 2018; Sikka & Bindra, 2010). Specific targeting the contact pathway may represent such a strategy. It is hoped that this approach may overcome functional obstacles in the quest for the ‘ideal anticoagulant’, that of treating or preventing thrombosis without affecting hemostasis. A key question in following up this approach is which of these factors is likely to be the more reliable target in the challenging endeavor of inhibiting blood clotting while maintaining hemostasis?

While physiological hemostasis depends primarily on the TF-factor pathway, the intrinsic pathway is not considered to be essential for hemostasis in vivo, even though its components appear to be involved in the pathogenesis of thrombosis (Doggen, Rosendaal, & Meijers, 2006; Meijers, Tekelenburg, Bouma, Bertina, & Rosendaal, 2000; Minnema, et al., 2000; Wheeler & Gailani, 2016). Epidemiologic data suggest that individuals with congenital factor XI deficiency while largely being protected against venous and arterial thrombosis (Rosen, Gailani, & Castellino, 2002; Salomon, Steinberg, Koren-Morag, Tanne, & Seligsohn, 2008; Salomon, et al., 2011; Tucker, et al., 2009), generally do not exhibit spontaneous bleeding (Duga & Salomon, 2013; James, Salomon, Mikovic, & Peyvandi, 2014). Additionally bleeding associated with severe deficiency, injury or surgery tends to be relatively mild, often not requiring treatment (Duga & Salomon, 2013; He, Chen, & He, 2012; James, et al., 2014; Preis, et al., 2017; Salomon, et al., 2008; Salomon, et al., 2011). A wide range of factor XI plasma levels has been found in the healthy population (Van Hylckama Vlieg, Callas, Cushman, Bertina, & Rosendaal, 2003) and elevated plasma levels of factor XI have been associated with increased risk of VTE and ischemic stroke (Folsom, et al., 2015; Meijers, et al., 2000; Salomon, et al., 2008; Siegerink, Rosendaal, & Algra, 2014; Yang, Flanders, Kim, & Rodgers, 2006). Conversely, lower factor XI levels have been associated with reduced risks of (recurrent) venous thrombosis and ischemic stroke, without evidence for an associated risk of major bleeding (Georgi, et al., 2019; Kyrle, Eischer, Šinkovec, & Eichinger, 2019).

Studies of patients with factor XII deficiency are limited. However, while factor XII deficiency does not cause abnormal bleeding (Kenne, et al., 2015), patients do not appear to be protected against thrombotic events (Key, 2014). In animal models both factor XI and XII inhibition results in protection from arterial and venous thrombosis without altering bleeding (Cai, et al., 2015; Larsson, et al., 2014; Matafonov, et al., 2014; Renne, et al., 2005; Rosen, et al., 2002; van Montfoort, et al., 2014; X. Wang, et al., 2005). These observations suggest that plasma factor XI level is a significant factor in thrombosis. Yet, therapeutic targeting of factor XI may also become relevant to interfere with vascular dysfunction, arterial hypertension (Kossmann, et al., 2017), and thrombus formation in atherosclerotic lesions (van Montfoort, et al., 2014), or during cardiopulmonary bypass (Pireaux, et al., 2019).

## 5. Shortcomings of current classes of anticoagulant therapeutics

While numerous approaches targeting intrinsic coagulation factors (XII, XI and IX) are currently under preclinical study (Al-Horani, 2020; Al-Horani & Afosah, 2018; Rai, Balters, & Agrawal, 2019; Weitz & Fredenburgh, 2017), only therapies utilizing small molecule inhibitors, antibodies and antisense oligonucleotides (ASO) targeting factors XII and XI have reached early phase human trials (DeLoughery, Olson, Puy, McCarty, & Shatzel, 2019). Here the use of an ASO to specifically reduce factor XI was the first of only two studies to complete phase II trials (Büller, et al., 2015; Weitz, et al., 2020). ASOs are relatively short, chemically modified single-stranded nucleic acid sequences that selectively pair to specific regions of mRNA resulting in endonucleolytic cleavage and degradation (Cosmi, 2016; DeVos & Miller, 2013; Heestermans & van Vlijmen, 2017; Kole, Krainer, & Altman, 2012) (Table 1). Currently over 60 ASO therapies are in, or have completed phase I/II trials (S. Zhu, Rooney, & Michlewski, 2020), with a substantial number of anti-thrombotic ASO therapeutics currently under development (Table 2). In animal studies, factor XI ASO can safely prevent thrombus formation without bleeding complications (J. R. Crosby, et al., 2013; van Montfoort, et al., 2014; Yau, et al., 2014; Younis, et al., 2012; H. Zhang, et al., 2010). A phase II study in surgical patients indicated that a specific factor XI ASO effectively protects patients against venous thrombosis with a relatively limited risk of bleeding (Büller, et al., 2015). However, this proof-of-concept trial was too small to assess the effect on other thrombotic end points. Additionally, changes in platelet counts in nonhuman primates treated with ASOs have been observed (Henry, et al., 2017), which has been attributed to peripheral clearance (Narayanan, et al., 2018) and could potentially impact hemostasis.

**Table 1.** General features of different next-generation therapeutics in the hemostatic system.

|  | **ASO/LNA/siRNA** | **miRNA** | **Antibody/biological** | **Small molecule** |
| --- | --- | --- | --- | --- |
| **Structure** | ASO/LNA: ~12-25 nucleotide single stranded RNA<br>siRNA: 21-23 nucleotide RNA duplex with 2 nucleotides 3'overhang | 19-25 nucleotide RNA duplex with 2 nucleotides 3'overhang. | Variable | Variable |
| **Targeting** | Fully complementary to mRNA | Partially complementary to mRNA, typically targeting the 3' untranslated region of mRNA | Binds catalytic domain, can mimic catalytic activities (bispecific antibodies) | Binds catalytic domain |
| **Mechanism of (gene) regulation** | Endonucleolytic cleavage and degradation of mRNA | Translational repression (particularly prevalent for secretory proteins), endonucleolytic cleavage and degradation of mRNA (especially in case of high complementarity between miRNA and mRNA) | Gain or loss of function by blocking catalytic activity of a enzyme or by chaperoning conformational complex formations (e.g. Emericizumab) | Blocks catalytic activity of enzyme |
| **Role, clinical application** | Therapeutic agent (loss of function) | Endogenous cell intrinsic gene regulatory mechanism<br>Diagnostic tool<br>Biomarker<br>Drug target and therapeutic agent (loss (mimic) and gain (antagomir) of function) | Therapeutic agent (loss and gain of function) | Therapeutic agent (loss of function) |
| **Advantages (therapeutic)** | Strong effect size<br>Easily synthesized in commercial laboratories | Mimics physiological gene regulation<br>Synergy with intrinsic target gene regulatory mechanisms<br>Easily synthesized in commercial laboratories | Blocks/mimics enzymatic activity | Blocks enzymatic activity |
| **Limitations (therapeutic)** | Unspecific effect on platelet count, others?<br>Potential off target effects<br>May induce partial IFN response and innate immune responses<br>Thrombocytopenia | Potential off target effects<br>Smaller effect size | Interference with diagnostics<br>Undesired immune reactions<br>Provokes production of inhibiting antibodies | Variable |
| **Therapeutic delivery** | Injection | Injection Exosomes? | Injection (s.c./i.v.) | Oral/injection |
| **Target (number)** | Single | Multiple | Single (multiple, bispecific antibodies) | Single |
| **Target** | Any gene (including 'undruggable' targets) | miRNA targeted genes (including 'undruggable' targets) | Enzymes | Enzymes |
| **Manufacturing process Development timeline** | Easily synthesized<br>Rapid | Easily synthesized<br>Rapid | Subject to purification and post-translational modifications issues<br>Slow | Unique chemical synthesis process for each compound<br>Slow |
| **Pharmacokinetics** | Consistent | Consistent | Individual | Individual |

**Table 2.** RNA based anti-thrombotic therapeutics.

| Status | Species | Type | Target | Outcome | References |
| --- | --- | --- | --- | --- | --- |
| Pre-clinical | Mouse | ASO | FII | Reduction of thrombosis, however deleterious effects on bleeding tendency | (J. Crosby, et al., 2009) |
| Pre-clinical | Mouse | ASO (ISIS 401025) | FII | Clotting times, PT and aPTT, significantly prolonged and correlated to degree of target inhibition | (Monia , et al., 2007) |
| Pre-clinical | Mouse | ASO | FMO3 | Impacted platelet responsiveness and rate of thrombus formation | (W. Zhu, et al., 2018) |
| Pre-clinical | Primate | ASO | FVII | Antithrombotic effects, however significant impairment of hemostasis | (Wallisch, et al., 2020) |
| Pre-clinical | Mouse | ASO | FVII | Prolongation of dilute PT, however no influence on dilute aPTT | (Vu, et al., 2015) |
| Phase 2 | Human | ASO (ISIS-FXIRx) | FXI | Reduced incidence and clot size of venous thromboembolism after surgery without increased bleeding risk | (Büller, et al., 2015) |
| Pre-clinical | Mouse | ASO | FXI | Safely prevented thrombus formation on acutely ruptured atherosclerotic plaques | (van Montfoort, et al., 2014) |
| Pre-clinical | Rabbit | ASO | FXI | Prolongation of time to catheter occlusion | (Yau, et al., 2014) |
| Pre-clinical | Primate | ASO | FXI | Antithrombotic effect without an increased risk of bleeding in arterial thrombosis model | (J. R. Crosby, et al., 2013) |
| Pre-clinical | Mouse | ASO | FXI | Reduction in thrombus formation in venous thrombosis model | (H. Zhang, et al., 2010) |
| Pre-clinical | Primate | ASO (ISIS 416858) | FXI | Increase in APTT with no effects on PT, platelets, or increased bleeding | (Younis, et al., 2012) |
| Pre-clinical | Rabbit | ASO | FXII | Prolongation of time to occlusion in a catheter thrombosis model | (Yau, et al., 2014) |
| Pre-clinical | Mouse | ASO | FXII | Reduction of arterial and venous thrombosis formation, without significant effect on normal hemostasis | (Revenko, et al., 2011) |
| Pre-clinical | Mouse | ASO | FXII | Prolongation of dilute aPTT, however no effect on the dilute PT | (Vu, et al., 2015) |
| Pre-clinical | Rabbit | ASO | FXII | Prolongation of time to catheter occlusion | (Yau, et al., 2014) |
| Pre-clinical | Rat | siRNA | FXII | Protection in arterial and venous thrombosis models | (Cai, et al., 2015) |
| Pre-clinical | Mouse | Antagomir | miR-148a | Reduction of thrombus formation following intravascular platelet activation via FcγRIIA | (Y. Zhou, et al., 2015) |
| Pre-clinical | Mouse | Mimic | miR-30c | Significant inhibition of arterial thrombosis formation | (Luo, et al., 2016) |
| Pre-clinical | Mouse | Mimic | miR-338-5p | Thrombus weight greatly decreased in inferior vena cava stenosis DVT model | (Y. Zhang, Zhang, et al., 2020) |
| Pre-clinical | Mouse | Antagomir | miR-374b-5p | Thrombus length and weight greatly decreased in inferior vena cava stenosis DVT model | (Y. Zhang, Miao, et al., 2020) |
| Pre-clinical | Mouse | Mimic | miR-495 | Reduction in occurrence and development of thrombosis of femoral vein | (Tang, et al., 2018) |
| Pre-clinical | Mouse | Mimic | miR-103a-3p | Thrombus size decreased in IVC ligation model | (S. Sun, et al., 2020) |
| Pre-clinical | Mouse | Antagomir | miR-19b-3p | Significant increase of antithrombin activity | (Nourse, et al., 2018) |
| Pre-clinical | Rat | ASO | PAI-1 | Protection against arterial thrombus formation | (Pawlowska, et al., 1998) |
| Pre-clinical | Rat | ASO<br>(MPO-16R) | PAI-1 | Significant delay in arterial thrombosis model occlusion time | (Pawlowska, et al., 2001) |
| Pre-clinical | Mouse | ASO | Prekallikrein | Reduction of arterial and venous thrombosis formation, without significant effect on normal hemostasis | (Revenko, et al., 2011) |
| Pre-clinical | Primate | ASO | Thrombopoietin | Sustained decline in platelet count | (Shirai, et al., 2019) |

RNA therapeutics offer the promise of uniquely targeting virtually any gene of interest involved in a particular disease with greater specificity, high potency, and decreased toxicity (Drosopoulos & Linardopoulos, 2013; Perwitasari, Bakre, Tompkins, & Tripp, 2013; Quemener, et al., 2020; S. Zhu, et al., 2020). The ASO-messenger(m)RNA heteroduplex silences gene expression by a variety of mechanisms including triggering RNase H activity resulting in mRNA degradation, steric blocking of ribosomal activity resulting in translational arrest, interfering with mRNA maturation by inhibiting splicing, destabilizing pre-mRNA in the nucleus or inhibiting 5’cap formation (J. H. Chan, Lim, & Wong, 2006; Crooke, 2000) (Table 1). While RNA therapeutic approaches have been applied to the development of new drugs and clinical trials currently being carried out (Tiemann & Rossi, 2009; Titze-de-Almeida, David, & Titze-de-Almeida, 2017), there are still concerns and challenges to be overcome. These include, but are not limited to, off-target effects (Snove & Holen, 2004; Yu, Jian, Yu, & Tu, 2019), triggering innate immune responses, efficacy and delivery into the target cell (X. Zhao, Pan, Holt, Lewis, & Lu, 2009). Recently, a new class of RNA therapeutic has attracted considerable attention, that of miRNA targeting.

## 6. miRNA targeting as novel class of therapeutic

MicroRNAs (miRNAs) are a class of short (17 to 25 bp), endogenous noncoding RNAs, that post-transcriptionally down-regulate target gene expression (Bartel, 2009; K. Chen & Rajewsky, 2007; Friedman, Farh, Burge, & Bartel, 2009). miRNAs are transcribed as primary transcripts, then processed in the nucleus by the microprocessor complex consisting of Drosha and DGCR8 to produce a pre-miRNA (S. Li & Patel, 2016). After further processing by Dicer (Krol, Loedige, & Filipowicz, 2010) the mature miRNA duplex is exported into the cytoplasm and incorporated into the RISC (Meister, 2013; Wilson & Doudna, 2013). This complex is guided by miRNA base pairing to a target gene mRNA resulting in translational inhibition and/or transcript degradation via one of several possible mechanisms (Fabian, Sonenberg, & Filipowicz, 2010; Oliveto, Mancino, Manfrini, & Biffo, 2017). In general miRNAs target mRNAs via the 3’ untranslated regions (3’UTR), while to a lesser extent, miRNAs can also carry out their inhibitory function by binding to the coding region or the 5’UTR of target mRNAs (Bartel, 2009).

Over 2600 human miRNAs have been identified (Kozomara, Birgaoanu, & Griffiths-Jones, 2019), regulating the majority of all human genes (Friedman, et al., 2009). Thus almost every biological process is modulated through miRNAs (Saliminejad, Khorram Khorshid, Soleymani Fard, & Ghaffari, 2019). Although miRNAs generally fine-tune gene expression (Mukherji, et al., 2011), they may also function as master regulators depending on the cellular environment (Lui, Jin, & Stevenson, 2015). Multiple miRNAs can cooperatively silence a single gene to gain regulatory specificity, with the targeting of particular network hub genes enabling the regulation of entire pathways (X. Li, et al., 2012). Additionally a single miRNA can target multiple genes (Bartel, 2009), allowing broad regulation of molecular networks (Megiorni, Cialfi, Dominici, Quattrucci, & Pizzuti, 2011). Perturbations of miRNA expression are observed in most disorders, with some of them even causally contributing to the development and progression of disease (Paul, et al., 2018; Saliminejad, et al., 2019). This underscores the extensive involvement of miRNAs in a variety of biological functions.

### 6.1 miRNA and therapy

The clear role of miRNAs in human disease has made them attractive targets for novel therapeutic intervention (Bajan & Hutvagner, 2020; Rupaimoole & Slack, 2017). miRNA-based therapeutics are actively being investigated and developed for clinical applications (Bonneau, Neveu, Kostantin, Tsongalis, & De Guire, 2019), with several currently the subject of clinical trials (Bajan & Hutvagner, 2020; Rupaimoole & Slack, 2017; Saliminejad, et al., 2019). Advances in RNA chemical modifications to improve stability, efficiency, target specificity and in vivo delivery while lowering toxicity have made miRNA-based therapeutics feasible (Egli & Manoharan, 2019; D. Lu & Thum, 2019). Several animal studies suggest that modified RNA-based oligonucleotides are specific, stable and non-toxic when delivered intravenously (Elmen, et al., 2008; Lanford, et al., 2010). Onset and duration of action are largely determined by the backbone chemistry of RNA-based oligonucleotides, with phosphorothioate-modified singlestranded ASOs exhibiting rapid transfer from blood into tissues (minutes to hours) and long halflives and prolonged activity (2-4 weeks) (Geary, Norris, Yu, & Bennett, 2015).

Depending on the desired outcome several different approaches to miRNA-based therapy may be taken. In situations where a miRNA is down-regulated in disease, the restoration of miRNA function can be achieved through the use of miRNA mimics or pre-miRNA. This involves RNA oligonucleotides, which match the corresponding miRNA sequence and can restore or increase miRNA function (Rupaimoole & Slack, 2017; van Rooij & Kauppinen, 2014). As an example, a miRNA mimic targeting miR-29, Remlarsen, is currently under Phase 2 trial in the prevention of fibrous scar tissue formation (Gallant-Behm, et al., 2019).

Where inhibition of miRNA activity is desired several strategies are available. Antagomirs are chemically modified synthetic antisense RNA oligonucleotides complementary to the target miRNA. An antagomir binds to and sequesters the endogenous target miRNA preventing it from binding the mRNA targets (Rupaimoole & Slack, 2017). One of the first miRNA therapies to be developed is Miravirsen, an antagomir against miR-122, which plays an important role in the replication cycle of the hepatitis C virus (Kaluzna, 2014). In human trials Miravirsen resulted in a decrease in viral RNA levels with no cytotoxicity (Janssen, et al., 2013; Ottosen, et al., 2015).

Although the use of such technology is widely developed for in vitro models, due to the ubiquitous nature of miRNA binding, application in vivo is hampered by undesired off-target effects (Y. Chen, Zhao, Tan, Zhang, & Fu, 2015; Messina, et al., 2016; Singh, Narang, & Mahato, 2011). Several approaches may assist in preventing off-target effects while maintaining the benefits of miRNA target diversity. The functional synergy of several miRNA mimics targeting a single gene, with each individually at sub-effective doses, allows for an increase in effect size and a reduction in side effects (Nourse, Braun, Lackner, Hüttelmaier, & Danckwardt, 2018). For miRNA inhibition a more precise alternative to inhibit miRNA function involves the use of an ASO designed to specifically bind an individual miRNA site within a target mRNA (termed target site blockers, or masks) preventing miRNAs from gaining access to the site (Louloupi & Orom, 2018; Z. Wang, 2011). This does not affect the expression of other genes thereby reducing potential off-target effects. Initial in vivo studies with miRNA targeting ASOs required relatively high (80 mg per kg) doses (Krützfeldt, et al., 2005), increasing the risk of off-target effects. The development of more efficient in vivo delivery systems that confer higher stability has reduced this (to the single-digit mg per kg range) (Forterre, Komuro, Aminova, & Harada, 2020). Although miRNA expression tends to exhibit tissue and cell type specificity (Nowakowski, et al., 2018; Sood, Krek, Zavolan, Macino, & Rajewsky, 2006), miRNA modulation in a target tissue may be accompanied by on-target side effects involving normal miRNA expression in non-target tissues, potentially leading to toxicity. To alleviate this miRNA delivery vehicles enabling tissue-specific targeting have been developed. Conjugates under investigation include antibodies (Cuellar, et al., 2015; Song, et al., 2005) and aptamers (McNamara, et al., 2006; J. Zhou & Rossi, 2017), while an alternative approach involves the production of targeted exosomes (Ohno, et al., 2013).

### 6.2 Comparison of miRNA to current therapeutic approaches

Protein expression (and hence the downstream functional consequences) depends not only on the rate of transcription, but on other regulatory mechanisms, including mRNA processing, stability, transport and translational regulation. These post-transcriptional mechanisms are principally mediated by the binding of RNA-binding proteins (RBPs), long non-coding RNAs (lncRNAs) and miRNAs to regulatory elements in the 3’UTRs of mRNAs (Figure 3). The emerging picture is that of complex and multifaceted layers of 3’UTR-mediated regulation, including competition and cooperativity of these regulatory components, to adjust protein synthesis in a spatiotemporal manner (Jens & Rajewsky, 2015; Tay, Rinn, & Pandolfi, 2014). Of note such mechanisms also tune the hemostatic system, for example in an acute event of blood loss to stockpile blood coagulation factors (Danckwardt, et al., 2011).

**Figure 2.**
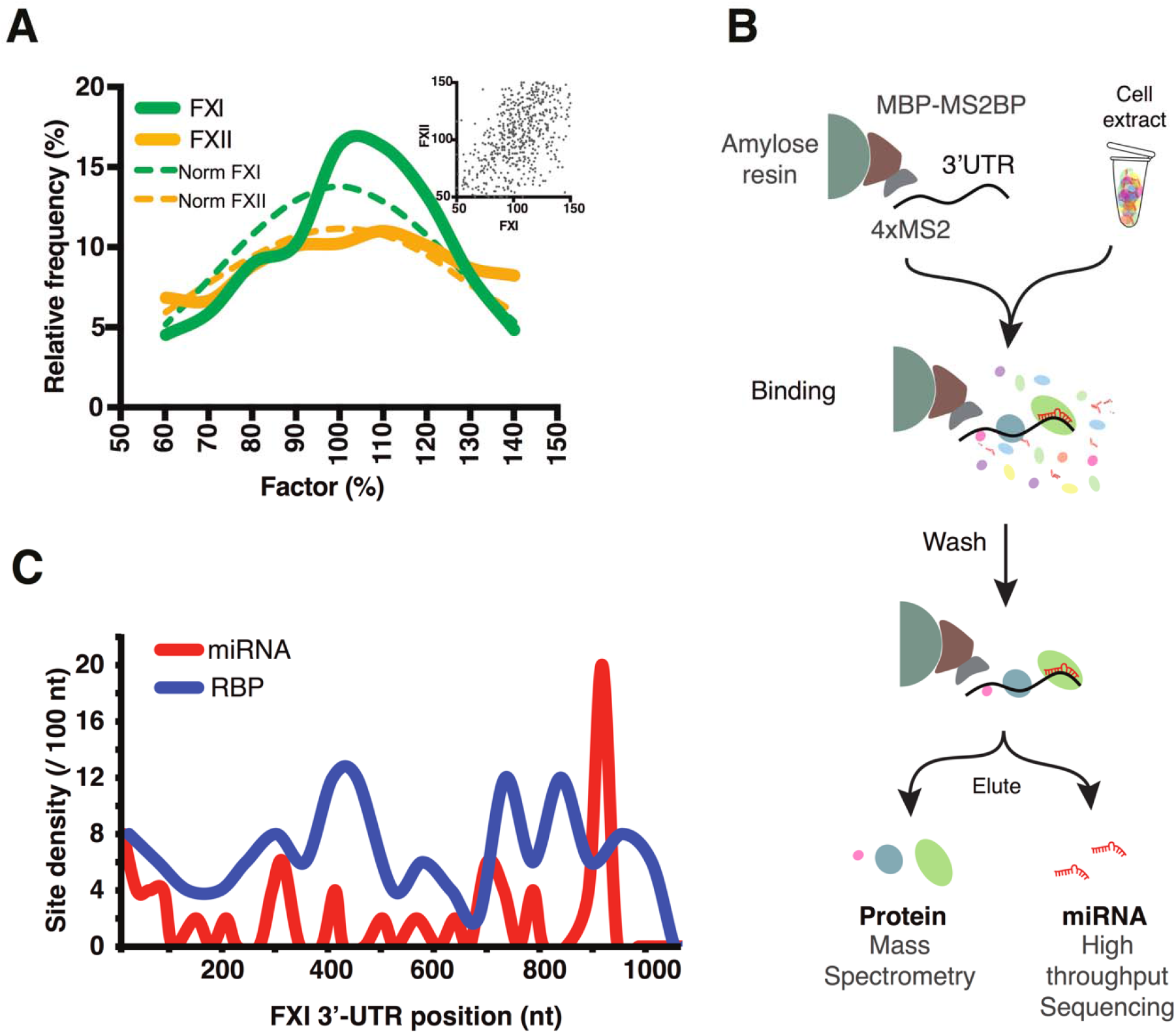
FXI is under tight post-transcriptional control. **(A)** FXI plasma levels exhibit a leptokurtic distribution as compared to platokurtic distribution type of FXII (levels of FXI and FXII from patients admitted to the University Medical Center Mainz from 2012 to 2018; values were limited to patients for which both FXI and FXII assays were performed and extremes (levels ?150% and ≤50%) have been excluded, normal distributions with standard deviations corresponding to those of FXI and FXII are shown dashed. Unpublished data). **(B)** Schematic of the in vitro pull down assay used to identify 3’UTR interacting miRNAs and RBP. Briefly baits consisting of in vitro transcribed 3’UTRs fused to copies of the bacteriophage MS2 stem-loop were immobilized by binding to amylose resin via recombinant MS2 coat protein-maltose binding (MBP-MS2BP) fusion protein. Immobilized baits were incubated with cell extracts to allow binding of 3’UTR interacting factors. Following washing protein–RNA complexes were eluted with maltose and either RNA or proteins were purified and identified using RNA-sequencing and mass spectrometry. **(C)** Density of potential sites for miRNA and RNA-binding proteins (RBPs) across FXI 3’UTR. 125 FXI-3’UTR/miRNA interactions were identified by miTRAP/RNA-seq (Nourse, et al., 2018) and of these 41 are mapped to the FXI 3’UTR using miRWalk target site prediction. 392 FXI-3’UTR/RBP interactions identified by miTRAP/MS (unpublished data) and of these 66 could be mapped to the FXI 3’UTR using RBPDB target site prediction. Site density calculated by number of sites present in 50 nt windows over length of the FXI-3’UTR.

**Figure 3.**
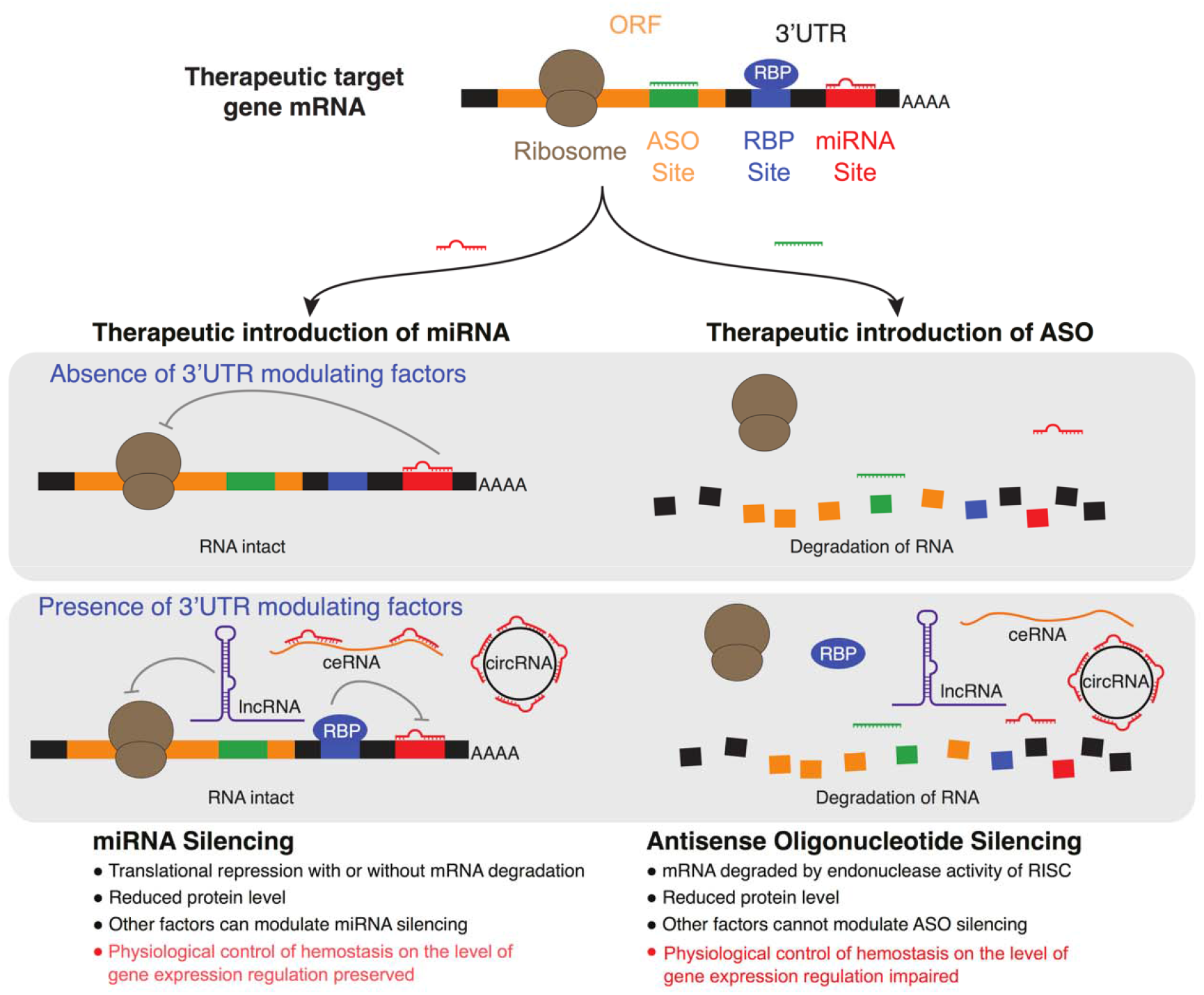
Function and consequences of therapeutic miRNA targeting versus antisense oligonucleotide (ASO) silencing. Post-transcriptional mechanisms mediated by the binding of RNA-binding proteins (RBPs), lncRNAs and miRNAs to regulatory elements in 3’UTRs of mRNAs, including competition and cooperativity of these regulatory components, to adjust protein synthesis in a spatiotemporal manner. As opposed to ASO silencing, and in many cases siRNAs, (right), which inevitably results in degradation of the target mRNA, the partial basepairing of miRNAs (left) prevents the cleavage activity of RISC with silencing predominately resulting from translational repression, and in some cases deadenylation, decapping and finally mRNA degradation (Bartel, 2009). Thus, miRNA-mediated therapeutic targeting without degradation of such target mRNAs preserves cell intrinsic regulatory mechanisms executed by 3’UTRs and their ‘interacting partners’ such as RBPs, miRNAs and/or other layers of control, such as ceRNA, circRNA miRNA sponges. Finally miRNA sites in proximity to RBP sites are often subject to mechanisms of physiological control (Danckwardt, et al., 2008; Jens & Rajewsky, 2015; Y. C. Lu, et al., 2014; Tay, et al., 2014), which may play important regulatory roles to control the delicate balance of hemostatic system in response to environmental cues (Danckwardt, et al., 2011).

As opposed to ASO silencing, and in many cases siRNAs, which exhibit perfect complementary to their targets, and inevitably result in degradation of the target mRNA (Figure 3 right, and Table 1) (Haley & Zamore, 2004; J. Martinez & Tuschl, 2004; Schwarz, et al., 2006), the partial base-pairing of miRNAs prevents the cleavage activity of RISC, with silencing predominately resulting from translational repression, and in some cases deadenylation, decapping and finally mRNA degradation (Fabian & Sonenberg, 2012; Huntzinger & Izaurralde, 2011) (Figure 3 left). Although the proportion of mRNA target degradation is reported to occur to a highly variable extent (Jonas & Izaurralde, 2015), a number of targets are almost exclusively repressed at the level of translation (Selbach, et al., 2008). How much each process contributes to down-regulation depends on characteristics, such as seed-flanking nucleotides, of the individual miRNA-mRNA pair (Grimson, et al., 2007; Nielsen, et al., 2007; Selbach, et al., 2008). Significantly, translational repression appears to be particularly relevant for secretory proteins (such as components of the hemostatic system) as miRNA translational repression is stronger for mRNAs produced at the endoplasmic-reticulum compared to free cytosolic ribosomes (Selbach, et al., 2008). Thus, miRNA-mediated therapeutic targeting without degradation of such target mRNAs preserves cell intrinsic regulatory mechanisms executed by 3’UTRs and their ‘interacting partners’ such as RBPs, miRNAs and/or other layers of control, such as ceRNA, circRNA miRNA sponges (Figure 3). Finally miRNA sites in proximity to RBP sites are often subject to mechanisms of physiological control (Danckwardt, Hentze, & Kulozik, 2008; Jens & Rajewsky, 2015; Y. C. Lu, et al., 2014; Tay, et al., 2014), which may play important regulatory roles to control the delicate balance of hemostatic system in response to environmental cues (Danckwardt, et al., 2011).

In the context of thrombosis and hemostasis, the versatility of miRNA-based approaches enables many therapeutic possibilities. miRNA mimics may be employed to silence procoagulant genes to treat thrombosis or alternatively, anticoagulant genes to treat bleeding. Conversely, antagomirs or target blockers may be used to relieve silencing of procoagulant genes to treat bleeding, or anticoagulant genes to treat thrombosis.

miRNA therapeutics are at an early stage of development and require time to reach the maturity point needed to reach the bedside (Beierlein, McNamee, & Ledley, 2017). One of the biggest challenges in the maturation of miRNA-based therapeutics is the identification of key miRNA candidates and targets. There is currently a relatively small number of experimentally validated miRNA:mRNA interactions (Lee, Kim, Muth, & Witwer, 2015) and here knowledge of the miRNA targetome in the hemostatic system is potentially a major trove for future targeted therapeutics (Nourse, et al., 2018), not only for the development of safe anticoagulants but also for developing treatments for coagulopathies.

### 6.3 Evidence for a functional role of miRNAs in the hemostatic system

Several lines of evidence implicate significant functional roles for miRNAs in the regulation of the hemostatic system. Firstly, a growing number of studies show essential contributions of miRNAs to the regulation of hemostatic (De Los Reyes-Garcia, et al., 2019; Jankowska, Sauna, & Atreya, 2020; Teruel-Montoya, Rosendaal, & Martinez, 2015) and thrombotic (Arroyo, et al., 2018; Hembrom, Srivastava, Garg, & Kumar, 2020; Jankowska, Sauna, et al., 2020; Morelli, Braekkan, & Hansen, 2020) functions (Table 3). Secondly, miRNAs have been demonstrated to directly regulate multiple hemostatic factors through interactions with the 3’UTR. This include key genes of the hemostatic system, including the fibrinogens FGA (Z. Chen, et al., 2010; Nourse, et al., 2018), FGB (Fort, et al., 2010) and FGG-alpha (Nourse, et al., 2018); the coagulation factors VII (Nourse, et al., 2018), VIII (Jankowska, Chattopadhyay, Sauna, & Atreya, 2020; Jankowska, McGill, et al., 2020; Nourse, et al., 2018; Sarachana, et al., 2015)], XI (Nourse, et al., 2018); TF (S. Li, Chen, et al., 2014; S. Li, Ren, et al., 2014; Sahu, et al., 2017; Teruel, et al., 2011; Witkowski, et al., 2016; X. Zhang, et al., 2011); prekallikrein (Nourse, et al., 2018); TFPI (Ali, et al., 2016; Arroyo, et al., 2017); antithrombin (SERPINC1) (Nourse, et al., 2018); protein c (Nourse, et al., 2018); protein z (Nourse, et al., 2018); protein z-dependent protease inhibitor (SERPINA10) (Nourse, et al., 2018); heparin cofactor 2 (SERPIND1) (Nourse, et al., 2018); VWF (L. Liu, et al., 2018); ADAMTS13 (L. Zhao, Hua, Li, Sun, & Wu, 2015); PAI-1 (SERPINE1) (Luo, et al., 2016; Marchand, Proust, Morange, Lompre, & Tregouet, 2012; Patel, Tahara, Malik, & Kalra, 2011); plasminogen (Nourse, et al., 2018) and the plasminogen activator PLAT (S. Li, et al., 2017). Additionally, miRNAs can impact hemostatic factors indirectly, for example fibrinogen via interleukin-6-mediated signaling (Brock, et al., 2011), factor IX by repressing nonsense-mediated mRNA decay (G. Wang, Chai, & Yang, 2016), PAI-1 via SMAD2 signaling (Liao, et al., 2014) and CXCL12 to reduce inflammatory response and thrombosis altering the expression of multiple factors including TF, PAI-1 and VWF (S. Sun, Chai, Zhang, & Lu, 2020).

**Table 3.** **A.** Functional miRNAs targeting the hemostatic system. The first two columns are based on the hemostatic miRNA targetome identified in Nourse et. al. (Nourse, et al., 2018) by miTRAP RNA pulldowns. The first column indicates interactions validated by functional miRNA-mimic-rescue luciferase assays, while the second column indicates interactions identified by miTRAP RNA pull-down alone (Nourse, et al., 2018). The last column indicates miRNAs identified from the literature to directly target the gene 3’UTR of blood coagulation factors indicated. **B.** Identity of functionally validated FXI targeting miRNAs and their role in cardiovascular disorders (AMI, acute myocardial infarction; VTE, venous thromboembolism; PAD, peripheral artery disease; CAD, coronary artery disease; DIC, disseminated intravascular coagulation).

| Gene | Gene Description | miRNAs identified in hemostatic miRNA targetome |  | Directly targeting miRNAs identified previously in literature |  |
| --- | --- | --- | --- | --- | --- |
|  |  | Directly targeting | Potential targeting | Directly targeting | References |
| **F11** | **Coagulation factor XI** | **10** | **126** | **4** | **(Salloum-Asfar, et al., 2014; Sennblad, et al., 2017; Vossen, et al., 2017)** |
| F8 | Coagulation factor VIII | 7 | 172 | 6 | (Jankowska, Chattopadhyay, et al., 2020; Jankowska, McGill, et al., 2020; Sarachana, et al., 2015) |
| SERPINA10 | Protein Z-Dependent Protease Inhibitor (ZPI) | 7 | 122 | 0 |  |
| SERPIND1 | Heparin co-factor II | 6 | 74 | 0 |  |
| PROZ | Protein Z | 6 | 57 | 0 |  |
| PLG | Plasminogen | 5 | 128 | 0 |  |
| FGG | Fibrinogen gamma chain | 4 | 127 | 0 |  |
| FGA | Fibrinogen alpha chain | 2 | 225 | 2 | (Z. Chen, et al., 2010; Fort, et al., 2010) |
| F7 | Coagulation factor VII | 2 | 103 | 0 |  |
| SERPINC1 | Antithrombin III | 2 | 57 | 0 |  |
| KLKB1 | Kallikrein B1 | 1 | 48 | 0 |  |
| FGB | Fibrinogen beta chain | 0 | 145 | 1 | (Fort, et al., 2010) |
| PROC | Protein C | 0 | 79 | 0 |  |
| PLAT | Plasminogen activator tissue type (tPA) | 0 | 76 | 2 | (Terao, et al., 2011; Yamashita, et al., 2015) |
| GP1BA | Glycoprotein Ib platelet alpha subunit | 0 | 0 | 2 | (Z. Zhang, Ran, Shaw, & Peng, 2016) |
| ADAMTS13 | ADAM metallopeptidase with thrombospondin type 1 motif 13 | 0 | 0 | 1 | (L. Zhao, et al., 2015) |
| **F12** | **Coagulation factor XII** | **0** | **0** | **0** |  |
| F13B | Coagulation factor XIII B polypeptide | 0 | 0 | 0 |  |
| F2 | Coagulation factor II (thrombin) | 0 | 0 | 0 |  |
| F5 | Coagulation factor V | 0 | 0 | 0 |  |
| HABP2 | Hyaluronan binding protein 2 | 0 | 0 | 0 |  |
| HRG | Histidine-rich glycoprotein | 0 | 0 | 0 |  |
| KNG1 | Kininogen 1 | 0 | 0 | 0 |  |
| SERPINF2 | Alpha-2-antiplasmin | 0 | 0 | 0 |  |

B
| miRNA | Thrombosis/Hemostasis associations | References |
| --- | --- | --- |
| **miR-103a-3p** | **Targets FXI** | (Nourse, et al., 2018) |
| miR-103a-3p | Involved in atherosclerosis and vascular inflammation by suppression of KLF4 | (Hartmann, et al., 2016) |
| miR-103a-3p | Biomarker for VTE | (Starikova, et al., 2015) |
| **miR-1255a** | **Targets FXI** | (Nourse, et al., 2018) |
| miR-1255a | Biomarker for stroke | (Edwardson, et al., 2018) |
| **miR-145-5p** | **Targets FXI** | (Sennblad, et al., 2017) |
| miR-145-5p | Biomarker for CAD, AMI, stroke, long-term outcome | (Dong, et al., 2015; Faccini, et al., 2017; Gao, et al., 2015; Jia, H. Wang, & Qu, 2015; Maitrias, et al., 2015; Meder, et al., 2011; Sal et al., 2014; M. Zhang, et al., 2017) |
| miR-145-5p | Impedes thrombus formation in atherosclerosis by targeting tissue factor and influencing platelet reactivity | (de Freitas, et al., 2017; Sahu, et al., 2017) |
| **miR-148b-3p** | **Targets FXI** | (Nourse, et al., 2018) |
| **miR-151a-3p** | **Targets FXI** | (Nourse, et al., 2018) |
| **miR-15b-5p** | **Targets FXI** | (Nourse, et al., 2018) |
| miR-15b-5p | Involved in development of hemophilic arthropathy | (Sen & Jayandharan, 2016) |
| miR-15b-5p | Biomarker for PAD | (Stather, et al., 2013) |
| miR-15b-5p | Influences platelet reactivity and clopidogrel response | (de Freitas, et al., 2017) |
| **miR-181a-5p** | **Targets FXI** | (Salloum-Asfar, et al., 2014; Sennblad, et al., 2017) |
| miR-181a-5p | Biomarker for AMI and PAD | (Stather, et al., 2013; J. Zhu, et al., 2016) |
| **miR-181b-5p** | **Targets FXI** | (Nourse, et al., 2018) |
| **miR-181b-5p** | Serum levels altered in patients with atherosclerosis | (X. Li & Cao, 2015) |
| **miR-191a-5p** | **Targets FXI** | (Salloum-Asfar, et al., 2014) |
| **miR-24-3p** | **Targets FXI** | (Nourse, et al., 2018) |
| miR-24-3p | Biomarker for acute cerebral infarction, arteriosclerosis obliterans, atherosclerosis and severe trauma | (L. J. Chen, et al., 2017; Zheng, Li, Liu, Qi, & Cao, 2018; J. Zhou Zhang, 2014; X. F. Zhu, et al., 2015) |
| **miR-30a-3p** | **Targets FXI** | (Nourse, et al., 2018) |
| miR-30a-3p | Biomarker for AMI and ischemic stroke | (Long, et al., 2012; Long, et al., 2013) |
| **miR-30d-3p** | **Targets FXI** | (Nourse, et al., 2018) |
| **miR-544a** | **Targets FXI** | (Vossen, et al., 2017) |
| **miR-96-5p** | **Targets FXI** | (Nourse, et al., 2018) |
| miR-96-5p | Biomarker for DVT and DIC | (Xie, et al., 2016; Y. Zhang, et al., 2014) |

A third line of evidence implicating miRNAs in the hemostatic system comes from the important roles miRNAs may play in the development of bleeding disorders and thrombosis. Blood miRNA levels are associated with many hemostatic diseases, suggesting not only an etiologic role, but their potential use as prognostic or diagnostic tools (J. Wang, Chen, & Sen, 2016). These include hemophilia A (Sarachana, et al., 2015), aberrant coagulation in sepsis (H. J. Wang, et al., 2014), venous thrombosis (Rodriguez-Rius, Lopez, Martinez-Perez, Souto, & Soria, 2020; X. Wang, et al., 2016; X. Wang, et al., 2019; Xiang, et al., 2019), venous thrombo-embolism (Hembrom, et al., 2020; Jiang, et al., 2017), arterial thrombosis (J. Lin, et al., 2016), acute pulmonary embolism (Guo, et al., 2014; Kessler, et al., 2016; Liu, Kang, & Liu, 2018; Q. Wang, et al., 2018; Xiao, et al., 2011; X. Zhou, et al., 2016), trauma-induced coagulopathy (L. J. Chen, Yang, Cheng, Xue, & Chen, 2017), coronary artery disease (Fichtlscherer, et al., 2010; Fujii, Sugiura, Dohi, & Ohte, 2016; X. Sun, Zhang, et al., 2012), atherosclerosis (Feinberg & Moore, 2016; Hosin, Prasad, Viiri, Davies, & Shalhoub, 2014; Menghini, Stohr, & Federici, 2014), stenosis (Dolz, et al., 2017; Jansen, et al., 2017), acute heart failure (Ovchinnikova, et al., 2016), congestive heart failure (Fukushima, Nakanishi, Nonogi, Goto, & Iwai, 2011; Tijsen, et al., 2010), ischemic stroke (Sorensen, Nygaard, & Christensen, 2016; Y. Wang, Ma, Kan, & Zhang, 2017) and systemic lupus erythematosus (SLE) (J. Q. Chen, et al., 2017; Kim, Jung, Jeon, Kim, & Suh, 2016; Perez-Sanchez, et al., 2016).

Cellular alterations in miRNAs are also associated with disease. In SLE and antiphospholipid syndrome patient monocytes miRNAs that target TF are down-regulated (Teruel, et al., 2011), while in patient neutrophils miRNAs linked to thrombosis and early atherosclerosis are expressed at lower levels (Perez-Sanchez, et al., 2016). Dysregulation of miRNA results in dysfunctional endothelial progenitor cells potentially trigger deep vein thrombosis (DVT) (Du, et al., 2019; Kong, et al., 2016; W. D. Li & Li, 2016; W. Wang, et al., 2019). On the other hand, polymorphisms affecting miRNA binding of hemostatic genes are associated with disease. Deletion of the miR-759 binding site of FGA is associated with the susceptibility to chronic thromboembolic pulmonary hypertension (Z. Chen, et al., 2010). SNPs in the 3’UTR of the F2, F8 and F11 genes are associated with increased plasma activity of the gene-products (Rosset, Vieira, Salzano, & Bandinelli, 2016; Sabater-Lleal, et al., 2012; Sennblad, et al., 2017; Voetsch & Loscalzo, 2004; Vossen, et al., 2017).

The importance of miRNAs in hemostasis is further highlighted by their role in platelet biology (De Los Reyes-Garcia, et al., 2019). Here miRNAs modulate the expression of target mRNAs important for hemostatic and thrombotic function (Dahiya, et al., 2015; Edelstein, et al., 2013; Elgheznawy, et al., 2015; Rowley, et al., 2016; Sunderland, et al., 2017). For example, miRNA levels are altered in platelets from patients with essential thrombocythemia and this is associated with elevated platelet counts and an increased risk of thromboembolic events (Tran, et al., 2020). Additionally in patient atherosclerotic plaques altered miRNA expression is observed (Raitoharju, et al., 2011).

Finally, miRNA treatment has been demonstrated to result in therapeutic response in thrombosis and hemostasis. In murine models of venous thrombosis, over-expression of miRNAs contributes to thrombus resolution (W. Wang, et al., 2019), reduces thrombogenesis (Sahu, et al., 2017), enhances endothelial progenitor cell migration and tubulogenic activity (Meng, et al., 2015) and significantly enhanced angiogenesis and thrombosis recanalization (L. L. Sun, et al., 2019). Antagomir inhibition has been shown to block miR-19b-3p-mediated silencing of SERPINC1 (antithrombin), resulting in an up-regulation of expression and increase in antithrombin activity, illustrating the in-principal druggability of the hemostatic system in a miRNA directed manner (Nourse, et al., 2018). Altogether multiple lines of evidence suggest a critical functional role of miRNA in the hemostatic system.

### 6.4 Factor XI is under significant miRNA control

Dysregulated miRNA-mediated control of factor XI expression can increase thrombotic risk. The F11 3’UTR SNP rs4253430 is associated with plasma factor XI levels and clotting time in idiopathic thrombophilia patients (Sabater-Lleal, et al., 2012; Sennblad, et al., 2017), and with increased factor XI activity and risk of VTE (Vossen, et al., 2017). This SNP disrupts a binding site for miR-544a preventing the miRNA from silencing factor XI expression and resulting in increased plasma factor XI activity (Vossen, et al., 2017).

The 3’UTR of the F11 mRNA is functionally targeted by miR-181a-5p (Salloum-Asfar, et al., 2014; Sennblad, et al., 2017). In healthy human liver samples a wider range of expression of F11 mRNA as compared with plasma levels was observed and the tissue levels were found to be inversely and significantly related to miR-181a-5p levels (Salloum-Asfar, et al., 2014). This suggests post-translational regulation may act to control plasma factor XI activity (Salloum-Asfar, et al., 2014). Another miR-181 family member, miR-181b-5p, also targets the 3’UTR of F11 mRNA (Nourse, et al., 2018) and members of this family are involved in several aspects of hemostasis including vascular inflammation and thrombogenicity (J. Lin, et al., 2016; X. Sun, Icli, et al., 2012; Witkowski, et al., 2020) and platelet activation (Dahiya & Atreya, 2020).

Other miRNAs demonstrated to functionally target the F11 3’UTR include miR-145, miR-103a-3p, miR-1255a, miR-148b-3p, miR-151a-3p, miR-15b-5p, miR-24-3p, miR-30a-3p, miR-30d-3p and miR-96-5p (Nourse, et al., 2018), with many of them exhibiting associations with vascular disease (L. J. Chen, et al., 2017; Cordes, et al., 2009; Elia, et al., 2009; Hergenreider, et al., 2012; Starikova, et al., 2015; J. Wang, et al., 2020; X. Wang, et al., 2019).

Finally, and most surprisingly in a recent endeavor to identify and characterize the hemostatic miRNA targetome (see further below) the largest quantity of functional miRNAs appear to target factor XI (Table 3).

The numerous roles played in hemostasis and thrombosis by miRNAs suggest the targeting of factor XI may be part of a network regulating coagulation, potentially offering new avenues for therapeutic targeting.

## 7. Factor XI miRNA as a novel anticoagulant therapy targeting avenue

In the search for novel and rationale therapeutic targets we recently performed a large-scale determination of functional miRNAs targeting the hemostatic system (Nourse, et al., 2018). Based on this study, and in conjunction with data presented above, we find factor XI to be the coagulation factor targeted by the largest number of miRNAs, while factor XII does not appear to be regulated by miRNAs (Table 3). Biologically, this finding is remarkable. It suggests that factor XI expression is under tight control via 3’UTR mediated regulation to ensure modest levels of factor XI protein while factor XII is not. This is in line with tighter regulation reflected by the leptokurtic distribution (i.e. narrower range) and lower percent coefficient variation (%CV) of factor XI plasma levels as compared to factor XII in our cohort (Figure 2A), which is consistent with earlier observations (Q. Chen, et al., 2015). The biological importance is corroborated by the functional consequences in patients where elevated factor XI levels predispose to thrombosis, while patients with FXI deficiency are protected from VTE and ischemic stroke (see above). Further support for the regulatory importance of the F11 gene 3’UTR comes from the observation that while the 3’UTR of F12 is relatively short (~150 nucleotides), that of F11 spans over ~1000 nucleotides, and as described previously F11 3’UTR SNPs are associated with plasma FXI activity and risk of venous thrombosis. Additionally, the F11 3’UTR is under control of alternative polyadenylation, which is consistent with increased regulation of gene expression and is associated with several diseases (Nourse, Spada, & Danckwardt, 2020). In addition to the high number of miRNAs, using RNA affinity pull-downs coupled to mass spectrometry, we identify here that the F11 3’UTR also binds numerous RNA binding proteins (RBPs) in close proximity to, and sometimes even in the direct target region of miRNAs (Figure 3B-C).

With regards to the important question as to which of the two components of the upstream intrinsic pathway (factor XI or factor XII) is likely to be a more reliable therapeutic target, our data suggests that the specific miRNA mediated control of factor XI may be a hitherto unrecognized rationale for the therapeutic targeting of factor XI. Additionally it may provide a novel avenue for this targeting.

It is important to realize that by mimicking the endogenous function of miRNAs to limit excessive expression, therapeutic miRNA delivery may possibly reflect a more physiological state in limiting factor XI levels to treat and prevent VTE while retaining sufficient hemostasis for the prevention of bleeding. Although miRNA targeting to silence factor XI could potentially have off-target effects, the functional synergy of several targeting miRNAs (at individually sub-functional levels) in a therapeutic cocktail may allow for an increase in effect size and a reduction in side effects (Nourse, et al., 2018). Finally and most importantly, as highlighted above (and see Table 1) miRNA-mediated targeting may also preserve cell intrinsic regulatory mechanisms, such as modulated ‘occlusion’ of miRNA binding sites by RNA binding proteins (RBPs) (Danckwardt, et al., 2008; Jens & Rajewsky, 2015; Y. C. Lu, et al., 2014), regulation through other noncoding RNAs or interaction with competing endogenous (ce)RNAs (i.e. miRNA sponges, circRNA) as well as other regulatory mechanisms (Tay, et al., 2014), ensuring the normal physiological fine tuning of the availability of the factor is maintained (Danckwardt, et al., 2011) (Figure 3, Table 1). Thus even in the presence of the miRNA therapeutic, RBP-mediated regulation (and other posttranscriptional gene regulatory mechanisms) may be in place to evade (therapeutic) miRNA-mediated silencing, for example in order to stockpile blood clotting factors in an acute event of blood loss (Danckwardt, et al., 2011).

Finally, although we have previously demonstrated the in-principle druggability of the hemostatic system in a miRNA-directed manner using intravenously administered LNA oligonucleotides (Nourse, et al., 2018), future therapies would ideally require a more convenient oral delivery route. This may be accomplished through novel delivery vehicles that allow oral miRNA uptake, for example milk exosomes (Manca, et al., 2018).

We thus believe that despite the impressive ‘juggernaut’ of anticoagulants, there are untapped therapeutic options that can help meet unmet medical needs. miRNA-based therapeutics, currently emerging as attractive therapies in many other disciplines (F. Wang, Zuroske, & Watts, 2020), may represent an avenue for correction of deregulated hemostasis and associated processes in the future. Given the plethora of miRNA-based therapeutic possibilities, such as miRNA mimicking, silencing, target specific miRNA target site blockers, knowledge of the miRNA targetome in the hemostatic system is potentially a major trove for future targeted therapeutics, not only for the development of safe anticoagulants but also for developing treatments for coagulopathies. In aid of these endeavors the first comprehensive hemostatic miRNA targetome is provided as a repository accompanying our recent paper (Nourse, et al., 2018).

## 8. Conclusion

RNA-based therapeutics have attracted considerable attention in the past decade because of their potential to treat numerous diseases, including cardiovascular disorders (D. Lu & Thum, 2019). Compared to conventional small therapeutic molecules, ASOs, siRNAs and miRNAs offer the advantage that they are highly effective and can act on “non-druggable” targets (e.g. proteins that lack an enzymatic function or whose conformation is not accessible to traditional drug molecules) as they can be designed to affect virtually any gene of interest (Daka & Peer, 2012). Innovative next-generation sequencing technologies provide platforms to assess the role of noncoding RNAs in the control of the hemostatic system and their utility as potential treatment targets (Nourse, et al., 2018).

Aligning with the more recent discovery of novel therapeutic targets in the hemostatic system (Büller, et al., 2015) factor XI appears to represent a potentially interesting target for miRNA-based approaches. Future studies are urgently needed to test this therapeutic principle in a clinical setting. This is particularly applicable in hemostaseology where monitoring of therapeutic effects remains a daily routine and thus assessment of the efficacy and safety of such therapeutic components could be simply implemented ushering in a novel therapeutic era with broad applicability.

## List of Abbreviations

aPTT: activated partial thromboplastin time
ASO: antisense oligonucleotide
ceRNA: competing endogenous RNA
circRNA: circular RNA
DOAC: direct oral anticoagulants
DVT: deep vein thrombosis
FGA: fibrinogen alpha
FGB: fibrinogen beta
FGG-alpha: fibrinogen gamma, alpha isoform
FII: factor 2
FIX: factor 9
FV: factor 5
FVIIa: factor 7 activated
FVIII: factor 8
FX: factor 10
FXI: factor 11
FXII: factor 2
miRNA: MicroRNA
RBP: RNA binding protein
RISC: RNA induced silencing complex
SNP: single nucleotide polymorphism
TF: tissue factor
UTR: untranslated regions
VKA: vitamin K antagonists
VSMC: smooth muscle of vascular walls
VT: venous thrombosis
VTE: venous thromboembolism
VTE: venous thromboembolism
VWF: Von Willebrand factor

## Acknowledgments

This work was supported by grants from the BMBF, DFG, DGKL and the Hella Bu□hler Prize for cancer research.

## Contribution

All authors contributed to the research and edited the manuscript; J.N. performed the data analysis; J.N. and S.D. wrote the manuscript.

